# Associations of ‘Relative corticosterone deficiency’ with genetic variation in *CYP17A1* and metabolic syndrome features

**DOI:** 10.1101/654269

**Authors:** Scott D Mackenzie, Andrew A Crawford, Daniel Ackermann, Katharina E Schraut, Caroline Hayward, Jennifer L Bolton, Christopher Saunders, Emad Al-Dujaili, Bernhard Dick, Geneviève Escher, Bruno Vogt, Menno Pruijm, Belen Ponte, James F Wilson, Mark W J Strachan, Jackie F Price, David I W Phillips, Scott M MacKenzie, Eleanor Davies, Rebecca M Reynolds, Brian R Walker

## Abstract

**Context and objective:** Common genetic variants in *CYP17A1* associate with higher blood pressure, putatively from impaired 17α-hydroxylase activity and mineralocorticoid excess. However, the same variants protect against obesity and insulin resistance. We tested whether *CYP17A1* variants that enhance 17α-hydroxylase activity cause ‘relative corticosterone deficiency’. Since corticosterone is thought to contribute disproportionately to negative feedback in the hypothalamic-pituitary-adrenal axis, we also tested whether lower corticosterone associates with higher cortisol and hence with metabolic syndrome.

**Design:** Cross-sectional studies within the population-based Orkney Complex Disease Study (ORCADES; n=2018), VIKING Health Study Shetland (VIKING; n=2098), East Hertfordshire study (EHERTS; n=279), Edinburgh Type 2 Diabetes Study (ET2DS; n=903), and the Swiss Kidney Project on Genes in Hypertension (SKIPOGH; n=888).

**Outcome measures:** Cortisol and corticosterone in morning plasma samples in ORCADES, VIKING and ET2DS, and in EHERTS in plasma following overnight dexamethasone suppression (0.25mg) and 30 mins after ACTH_1-24_ (1µg); cortisol and corticosterone metabolites in day and night urine samples in SKIPOGH. Features of the metabolic syndrome including body mass index, systolic blood pressure, lipid profile, fasting glucose, fasting insulin and HOMA-IR.

**Results:** In ORCADES, ET2DS and SKIPOGH, *CYP17A1* variants were associated with corticosterone:cortisol ratio. In ORCADES, VIKING and ET2DS there were consistent associations of morning plasma cortisol and corticosterone with BMI, blood pressure, lipid profile, fasting glucose and HOMA-IR. In EHERTS, however, after dexamethasone suppression and ACTH_1-24_ stimulation, impaired glucose tolerance and insulin sensitivity were associated with higher cortisol but lower corticosterone. Similarly, in SKIPOGH, low corticosterone:cortisol metabolite ratios were associated with high BMI and dyslipidemia.

**Conclusions:** ‘Relative corticosterone deficiency’, due to a primary alteration in adrenal steroidogenesis favouring cortisol over corticosterone, may mediate the associations of genetic variation in *CYP17A1* with metabolic syndrome. However, additional determinants of variation in plasma corticosterone are likely to explain its generally positive associations with features of metabolic syndrome.

## INTRODUCTION

In genome-wide association studies (GWAS) common variants in the *CYP17A1* locus are consistently associated with hypertension (1–4) and cardiovascular disease (5, 6). *CYP17A1* encodes the steroidogenic enzyme CYP17A1, which is expressed in human but not rat or mouse adrenal cortex and catalyses 17α-hydroxylase activity, converting precursors for synthesis of 11-deoxycorticosterone and corticosterone into 17-hydroxylated precursors for synthesis of 11-deoxycortisol and cortisol. Rare mutations causing near-absent CYP17A1 activity result in low-renin hypertension (7), attributed to accumulation of the mineralocorticoid 11-deoxycorticosterone. Ratios of steroid metabolites reflecting CYP17A1 activity are highly heritable (8). Associations of more common variation in *CYP17A1* with hypertension might therefore be explained by reduced 17α-hydroxylase activity.

Intriguingly, *CYP17A1* risk alleles for hypertension are also associated with lower body mass index (BMI) (9–11) and enhanced insulin sensitivity (12). Conversely, alleles which are protective for hypertension, and therefore predicted to increase 17α-hydroxylase activity, confer increased risk of obesity and insulin resistance. This paradox is unexplained, but may relate to variations in production of the major glucocorticoid, cortisol. Elevated cortisol in Cushing’s syndrome causes obesity and insulin resistance, while plasma cortisol is more subtly increased in metabolic syndrome (13). However, if elevated 17α-hydroxylase activity contributes to obesity and insulin resistance by increasing cortisol production, why would plasma cortisol concentration not be ‘corrected’ by enhanced negative feedback suppression of ACTH secretion and a compensatory fall in cortisol production? An explanation may be found by considering the role of the second glucocorticoid in humans, corticosterone.

Widely neglected in humans, corticosterone circulates at concentrations of 5-10 % those of cortisol. However, concentrations of corticosterone in human cerebrospinal fluid and brain are relatively high, at ∼30% of cortisol (14, 15). This discrepancy has been attributed to differential trans-membrane trafficking of steroids by the ATP-binding cassette (ABC) transporter ABCB1, which is expressed in the human CNS and selectively exports cortisol rather than corticosterone (15). Corticosterone may therefore make a disproportionate contribution to negative feedback control of the hypothalamic-pituitary-adrenal (HPA) axis within the CNS. Notably, the endogenous negative feedback signal is known to be impaired in people with metabolic syndrome (16).

We hypothesised that individuals with a genetically determined increase in 17α-hydroxylase activity have relatively low corticosterone which causes impaired negative feedback to the HPA axis, and hence sustains higher plasma cortisol. We predict this pattern of ‘relative corticosterone deficiency’ is associated with obesity and insulin resistance but lower blood pressure. We tested this hypothesis in population-based cohorts, by studying the associations of corticosterone and cortisol in plasma and of their metabolites in urine with: (i) genetic variation in *CYP17A1*; and/or (ii) features of metabolic syndrome.

## MATERIALS AND METHODS

### Participants

All studies conformed with the Declaration of Helsinki and ethical approval and written informed consent were obtained.

#### Orkney Complex Disease Study (ORCADES)

The ORCADES study is a genetic epidemiological study based in the Scottish archipelago of Orkney, comprising 2039 subjects aged 18-100 years, with at least two Orcadian grandparents, recruited between 2005 and 2011 (17). Subjects attended a local or mobile venepuncture clinic between 0730h and 1100h (mean 0923 h ± SD 46 min), after fasting from 2200h the previous night. On another occasion subjects attended for measurement of weight, height and blood pressure. Genotyping was undertaken as described (17).

#### Viking Health Study – Shetland (VIKING)

The VIKING study is a population cohort study based in an isolated population in the north of Scotland that recruited 2105 volunteers between 2013 and 2015. Subjects were required to have at least two grandparents from the Shetlands in order to participate. Each participant attended a measurement clinic and a venepuncture clinic to give a fasting blood sample. Participants’ DNA was genotyped using the Illumina HumanOmniExpressExome8v1-2_A.

#### Edinburgh Type 2 Diabetes Study (ET2DS)

The Edinburgh Type 2 Diabetes Study (ET2DS) is a prospective cohort study comprising 1066 men and women with type 2 diabetes, living in Lothian, Scotland. Recruitment and study design has been reported (18). Briefly, participants aged 60-75 years with a diagnosis of type 2 diabetes according to WHO criteria (19) were recruited from a clinical database. Following overnight fast, subjects attended a research clinic at 0800-0830h where they underwent venepuncture and physical examination. Genotyping was performed by KBioscience (Herts, UK) using a competitive allele-specific PCR system (KASPar).

#### East Hertfordshire Study (EHERTS)

All births in EHERTS from 1911 onwards were notified by the attending midwife (20). As described, these records were used to recruit 309 women and 370 men, with no history of diabetes, for a standard 75-g oral glucose tolerance test (OGTT), performed at a clinic at 0800-1020 h following overnight fast from 2100 h (21). In follow up studies (21), 312 of these individuals, aged 67-78 years, ingested 0.25 mg dexamethasone at 2200 h, fasted overnight and attended a local clinic at 0830h next morning. A baseline blood sample was obtained before 1.0µg ACTH_1-24_ (tetracosactrin, Synacthen, Alliance, Chippenham, UK) was administered intravenously, and repeat samples obtained after 30 minutes.

#### Swiss Kidney Project on Genes in Hypertension (SKIPOGH)

The SKIPOGH study is a family-based cross-sectional study comprising 1093 subjects aged 18-82 years from 2 regions (Bern and Geneva) and 1 city (Lausanne) of Switzerland (8). A random sample of the inhabitants were invited to participate if they were of European ancestry and had at least 1 first degree family member also willing to participate. They attended hospital in the morning after an overnight fast for venepuncture and physical examination. Five consecutive blood pressure (BP) measurements were taken from the arm with the higher BP, and the average of the last four was used. Urine samples were collected separately for day- and night-time periods, as described (8). DNA samples were genotyped on the Illumina Metabochip array.

### Laboratory analyses

In ORCADES, VIKING, EHERTS, and SKIPOGH, the homeostasis model assessment was used to quantify insulin resistance (HOMA-IR) (22). Serum cortisol was measured by radioimmunoassay with Guildhay antisera (23). After exclusion of individuals prescribed glucocorticoid therapy within the previous 3 months (ORCADES n = 21; ETD2S n=163; EHERTS n = 33), there was sufficient sample for analysis of corticosterone in 2018 individuals from ORCADES, 2098 in VIKING, 903 in ET2DS, and 279 in EHERTS. An in-house radioimmunoassay, modified for microtiter plate scintillation proximity assay (SPA), was used to measure plasma corticosterone with assay characteristics as previously reported (24); cross-reactivity for dexamethasone and cortisol was less than 1%. Results were accepted if the coefficient of variation between duplicates was <15%. In SKIPOGH, urinary steroid metabolites were extracted and analysed by gas chromatography-mass spectrometry (GC-MS) as previously described (8); analyses comprised 888 individuals after exclusion of 205 subjects with missing steroid metabolites. To estimate the apparent CYP17A1 activity, the ratio (tetrahydro-11-dehydrocorticosterone (THA) + tetrahydrocorticosterone (THB) + 5α-tetrahydrocorticosterone (5αTHB)) / (tetrahydrocortisone (THE) + tetrahydrocortisol (THF) + 5α-tetrahydrocortisol (5αTHF)) was calculated.

### Statistical analysis

Associations with genotype were analysed using Stata v14.1 (Stata Statistical Software: Release 14. College Station, TX: StataCorp LP). Given limited statistical power, we restricted analysis to candidate single nucleotide polymorphisms (SNPs) in *CYP17A1* which have functional effects; these included rs2486758, located in the *CYP17A1* promoter region and associated with ovarian steroidogenesis (25, 26), and the other SNPs in Table 2 which affect CYP17A1 transcription *in vitro* (27). All analyses were adjusted for age and sex. Because clearance of glucocorticoids is increased and plasma cortisol decreased in obesity (28), relevant analyses were adjusted for BMI. Because timing of sampling varied in ORCADES, venesection time (as minutes after the first sample in the cohort) was included as a predictor variable in all models (29). In ORCADES many of the participants are related; analysis of associations of SNPs with plasma glucocorticoid concentrations and metabolic risk factors was therefore undertaken following adjustment after fitting both the first 3 principal components of ancestry and the kinship matrix using a mixed model (mmscore function of GenABEL for the association test in ProbABEL) (30) under an additive model. The kinship matrix used the identity-by-state function of GenABEL (using weight = “freq” option).

Other analyses for ORCADES, VIKING, ET2DS and EHERTS were undertaken using Minitab (version 16; State College PA). Continuous response variables were normalised by log transformation. Student’s t-tests were used for group comparisons of corticosterone:cortisol ratios and responses to ACTH. Simple linear regression analysis of corticosterone or cortisol with a given ‘response’ (or dependent) variable was used to determine unadjusted p values for the regression co-efficient. Analyses were repeated with potentially confounding co-variables in multiple regression analyses, adjusting for age, sex, BMI and time of sampling. Histograms and normal plots of residuals were examined to confirm the validity of the linear regression model. To maintain this validity, and to minimise confounding due to prescribed medication, those prescribed glucose-lowering medications (see Table 1 for numbers) were excluded from regression analyses involving glucose or insulin in the ORCADES, VIKING and EHERTS cohorts. Because the majority of participants in ET2DS were prescribed these agents, their prescription was encoded as a binary variable and entered into the regression equation. This binary variable approach was also used in all cohorts to adjust for the effect of relevant medications on lipids and blood pressure.

**Table 1.** Characteristics of participants

|  | Cohort |  |  |  |  |
| --- | --- | --- | --- | --- | --- |
|  | ORCADES | EHERTS | ET2DS | SKIPOGH | VIKING |
| Sex (M/F) | 794/1224 (39/61%) | 189/90 (68/32%) | 476/427 (53/47%) | 380/508 (43/57%) | 838/1260 (40/60%) |
| Age (years) | 53.3 ± 0.3 (range 16-91) | 71.3 ± 0.2 (range 67-78) | 67.9 ± 0.1 (range 60-68) | 47.1 ± 0.6 (range 18-82) | 49.8 ± 0.3 (range 18-92) |
| BMI (kg/m<sup>2</sup>) | 27.7 ± 0.1 | 27.1 ± 0.2 | 31.2 ± 0.2 | 24.7 ± 0.1 | 27.4 ± 0.1 |
| Taking anti-hypertensive (n (%)) | 412 (20%) | 105 (38%) | 704 (79%) | 124 (14%) | 344 (16%) |
| Systolic BP (mmHg) | 129.9 ± 0.4 | 159.3 ± 1.3 | 133.3 ± 0.6 | 116.8 ± 0.6 | 127.0 ± 0.4 |
| Diastolic BP (mmHg) | 75.5 ± 0.2 | 86.9 ± 0.7 | 69.2 ± 0.3 | 75.1 ± 0.3 | 74.7 ± 0.2 |
| Plasma corticosterone at baseline (nM) | 50.7 ± 1.2 | 20.2 ± 1.0* | 22.3 ± 0.5 | n/a | 20.3 ± 1.4 |
| Plasma corticosterone 30 min after ACTH<sub>1-24</sub> (nM) | n/a | 84.0 ± 2.5 | n/a | n/a | n/a |
| Plasma cortisol at baseline (nM) | 724 ± 7 | 188 ± 6* | 732 ± 6 | n/a | 293 ± 4 |
| Plasma cortisol 30 min after ACTH<sub>1-24</sub> (nM) | n/a | 460 ± 8 | n/a | n/a | n/a |
| Lipid lowering therapy (n (%)) | 255 (13%) | 71 (25%) | 755 (84%) | 34 (3.8%) | n/a |
| Total cholesterol (mM) | 5.39 ± 0.03 | 6.82 ± 0.07 | 4.28 ± 0.03 | 5.1 ± 0.03 | 5.30 ± 0.02 |
| LDL cholesterol (mM) | 3.37 ± 0.02 | 4.89 ± 0.08 | n/a | 3.11 ± 0.03 | 3.33 ± 0.02 |
| HDL cholesterol (mM) | 1.50 ± 0.01 | 1.33 ± 0.02 | 1.28 ± 0.01 | 1.55 ± 0.01 | 1.52 ± 0.01 |
| Triglycerides (mM) | 1.15 ± 0.02 | 1.54 ± 0.05 | n/a | 1.02 ± 0.02 | 1.00 ± 0.01 |
| Medication for diabetes (n) | 47 (2.3%) | 10 (3.6%) | 734 (81%) | 8 (0.8%) | 44 (2.1%) |
| Oral agents |  |  | 660 (74%) | 7 (0.7%) |  |
| Insulin |  |  | 156 (18%) | 1 (0.1%) |  |
| Fasting plasma glucose (mM) | 5.33 ± 0.02 | 5.99 ± 0.08 | 7.57 ± 0.07 | 5.14 ± 0.02 | 4.88 ± 0.02 |
| Plasma glucose 30 min after oral glucose (mM) | n/a | 9.38 ± 0.14 | n/a | n/a | n/a |
| Plasma glucose 120 min after oral glucose (mM) | n/a | 7.13 ± 0.18 | n/a | n/a | n/a |
| Fasting plasma insulin mU/L | 6.44 ± 0.10 | 7.18 ± 0.35 | n/a | 6.19 ± 0.21 | 8.31 ± 0.80 |
| HOMA-IR | 1.59 ± 0.34 | 1.96 ± 0.11 | n/a | 1.48 ± 0.06 | 1.86 ± 0.17 |
| **Daytime urinary steroids** |  |  |  |  |  |
| Tetrahydrodehydrocorticosterone (µg/day) | n/a | n/a | n/a | 67.3 ± 1.3 | n/a |
| Tetrahydrocorticosterone (µg/day) | n/a | n/a | n/a | 91.7 ± 1.7 | n/a |
| 5α-Tetrahydrocorticosterone ($\mu\text{g/day}$) | n/a | n/a | n/a | 188.5 ± 4.2 | n/a |
| Tetrahydrocortisol ($\mu\text{g/day}$) | n/a | n/a | n/a | 1080.5 ± 18.2 | n/a |
| 5α-Tetrahydrocortisol ($\mu\text{g/day}$) | n/a | n/a | n/a | 779.9 ± 20.8 | n/a |
| Tetrahydrocortisone ($\mu\text{g/day}$) | n/a | n/a | n/a | 1861.2 ± 33.8 | n/a |
| **Nighttime urinary steroids** |  |  |  |  |  |
| Tetrahydrodehydrocorticosterone ($\mu\text{g/night}$) | n/a | n/a | n/a | 27.3 ± 0.7 | n/a |
| Tetrahydrocorticosterone ($\mu\text{g/night}$) | n/a | n/a | n/a | 41.8 ± 1 | n/a |
| 5α-Tetrahydrocorticosterone ($\mu\text{g/night}$) | n/a | n/a | n/a | 60.7 ± 1.7 | n/a |
| Tetrahydrocortisol ($\mu\text{g/night}$) | n/a | n/a | n/a | 362.9 ± 8.2 | n/a |
| 5α-Tetrahydrocortisol ($\mu\text{g/night}$) | n/a | n/a | n/a | 247.1 ± 6.8 | n/a |
| Tetrahydrocortisone ($\mu\text{g/night}$) | n/a | n/a | n/a | 642.6 ± 14.2 | n/a |

Data presented as mean ± SEM. n/a = not available.
\* Dexamethasone suppressed in EHRTS cohort

The magnitude of the association of plasma cortisol and corticosterone with response variables is presented using standardised Z scores of log transformed variables in the adjusted regression models. The coefficients from these regression models are interpreted as the SD change in log transformed outcome for a 1SD higher log transformed cortisol or corticosterone. This allows direct comparison of all associations investigated.

In SKIPOGH statistical analyses were conducted using STATA 14.0 (StataCorp, College Station, Texas, USA). Simple mixed linear or multiple regression analyses were used to explore associations of total urinary corticosterone (THA+THB+5α-THB) or cortisol (THE+THF+5α-THF) metabolites, while taking familial correlations into account using a random family-effect. In the multiple regression model, analyses were adjusted for age, sex, study center, BMI (unless BMI was the outcome variable) and relevant medication. (THA+THB+5αTHB)/(THE+THF+5αTHF) ratio (i.e. CYP17A1 ratio) was analysed after log transformation in a mixed linear model to investigate associations with variables of interest. Covariates included age, sex, study center, and antihypertensive, lipid- and glucose-lowering treatment, BMI (unless BMI was the outcome variable), estimated GFR, urine flow rate, urinary sodium, potassium and creatinine excretion (24h per kg body weight).

## RESULTS

Participant characteristics are summarised in Table 1. Plasma cortisol and corticosterone were positively correlated in ORCADES and ET2DS cohorts (Figure 1A), in VIKING (Coefficient correlation = 0.08, p<0.001) and in both dexamethasone-suppressed and ACTH-stimulated samples in EHERTS (Figure 1B), while urinary cortisol and corticosterone metabolites were positively correlated in SKIPOGH (Figure 1C and 1D). After ACTH stimulation in EHERTS, the increase from baseline was greater for corticosterone than cortisol (6.2±0.3 vs 3.9±0.4 fold, respectively; p<0.001).

**Figure 1.**
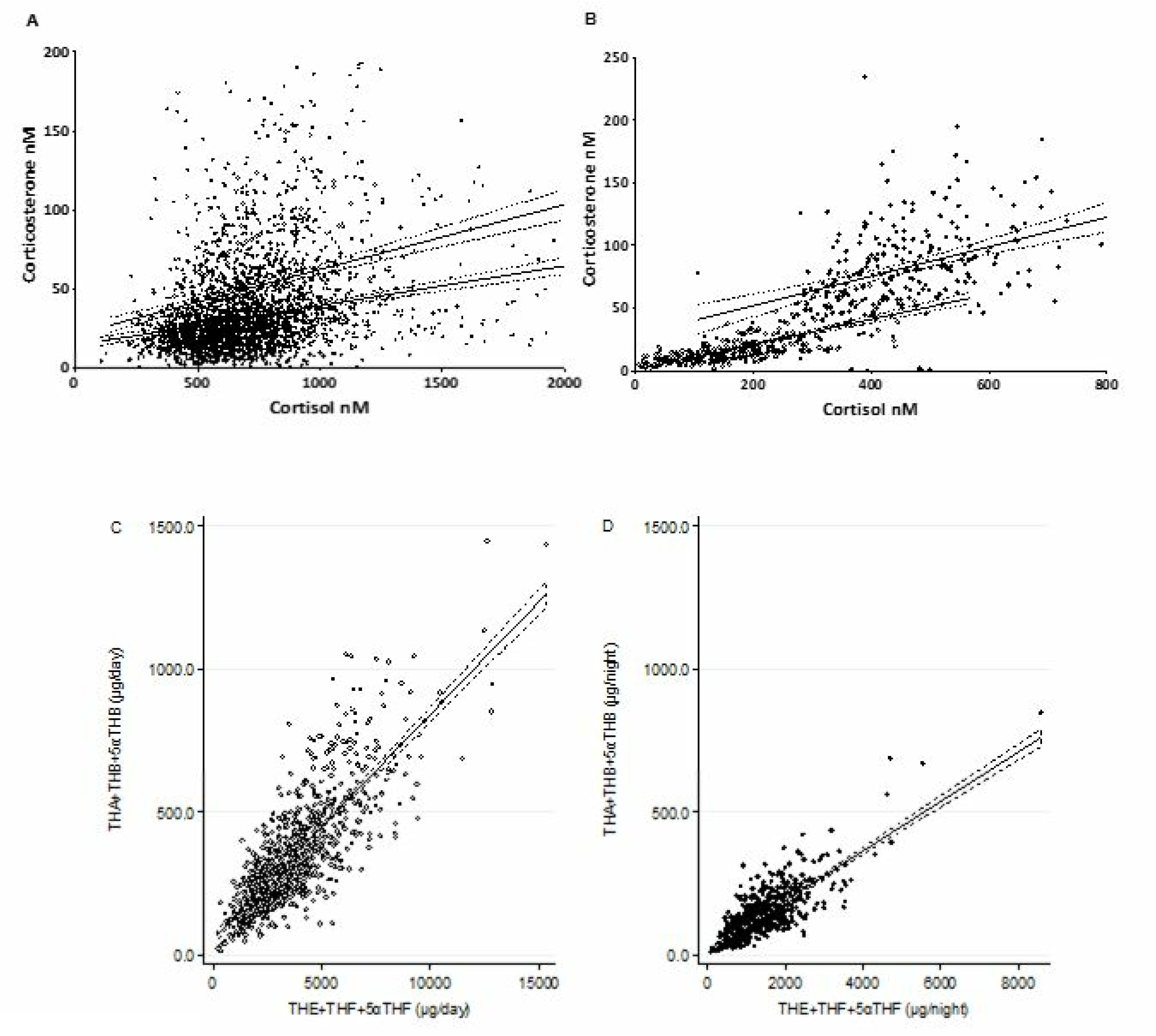
Relationships between plasma cortisol and corticosterone. A) Plasma cortisol and corticosterone in the Orkney complex diseases study group (ORCADES, open symbols) and the Edinburgh Type 2 Diabetes Study (ET2DS, closed symbols). Lines indicate simple linear regression with 95% confidence bands. Coefficient of determination (r^2^): ORCADES = 0.062, p<0.001; ET2DS = 0.070, p<0.001. B) Cortisol and corticosterone in the East Hertfordshire Cohort. Sampling was undertaken following overnight dexamethasone (0.25mg) suppression testing (pre synacthen, open symbols); and 30 minutes following intravenous injection of ACTH_1-24_ (synacthen 1µg, closed symbols). Lines indicate simple linear regression with 95% confidence bands. Coefficient of determination (r^2^): pre-synacthen = 0.488, p<0.001; post synacthen = 0.168, p<0.001. C and D) Total urinary cortisol and corticosterone metabolites in the Swiss Kidney Project on Genes in Hypertension. Sampling was undertaken for day (C, open symbols) and night (D, closed symbols) separately. Lines indicate simple linear regression with 95% confidence bands. Coefficient of determination (r^2^): day = 0.641, p<0.001; night = 0.69, p<0.001.

### Associations of plasma corticosterone and cortisol with *CYP17A1* genotype

In ORCADES, associations of functional SNPs in *CYP17A1* with morning plasma cortisol and corticosterone are shown in Table 2. The minor allele (C; frequency 0.16) of rs2486758 was associated with higher plasma corticosterone and corticosterone:cortisol ratio but not with plasma cortisol. These associations were replicated in ET2DS, in which the minor allele (C; frequency 0.21) of rs2486758 similarly tended to be associated with higher plasma corticosterone (β=0.074±0.043, p=0.09) and corticosterone:cortisol ratio (β=0.078±0.040, p=0.05) but not with cortisol (β=-0.004±0.017, p=0.83). Similar associations were observed in SKIPOGH with the minor allele (C; frequency 0.23) of rs2486758 associated with higher overnight urinary corticosterone (β=11.0±5.7, p=0.05) and with day and night urinary corticosterone:cortisol ratio (β=0.004±0.002, p=0.03; β=0.004±0.002, p=0.05), but not with daytime or overnight urinary cortisol (β=-133.5±96.6, p=0.17; β=11.1±44.6, p=0.80). There was no evidence of an association between *CYP17A1* variants with morning plasma cortisol or corticosterone in the VIKING cohort.

**Table 2.** Associations of functionally significant SNPs in *CYP17A1* with morning plasma cortisol and corticosterone

| Study | SNP ID | Minor allele frequency | Cortisol |  | Corticosterone |  | Corticosterone:cortisol ratio |  |
| --- | --- | --- | --- | --- | --- | --- | --- | --- |
| | | | $\beta$ | P | $\beta$ | p | $\beta$ | p |
| ORCADES | rs2486758 | 0.16 | 0.000 | 0.98 | 0.074 | 0.01 | 0.070 | 0.01 |
|  | rs138009835 | 0.09 | 0.003 | 0.89 | 0.004 | 0.91 | 0.003 | 0.93 |
|  | rs2150927 | 0.36 | -0.008 | 0.52 | 0.023 | 0.29 | 0.032 | 0.14 |
| ET2D | rs2486758 | 0.21 | -0.004 | 0.83 | 0.074 | 0.09 | 0.077 | 0.05 |
|  | rs138009835 | 0.08 | 0.006 | 0.82 | 0.004 | 0.95 | -0.002 | 0.97 |
|  | rs2150927 | 0.36 | -0.005 | 0.70 | -0.025 | 0.49 | -0.019 | 0.56 |
| VIKING | rs2486758 | 0.21 | -0.003 | 0.85 | -0.009 | 0.77 | -0.004 | 0.88 |
|  | rs138009835 | 0.09 | 0.006 | 0.78 | -0.024 | 0.60 | 0.040 | 0.26 |
|  | rs2150927 | 0.35 | 0.012 | 0.35 | 0.019 | 0.51 | -0.012 | 0.58 |
| SKIPOGH (day) | rs2486758 | 0.23 | -133.5 | 0.17 | 6.86 | 0.60 | 0.004 | 0.03 |
|  | rs138009835 | 0.10 | -119.7 | 0.37 | -27.58 | 0.14 | -0.004 | 0.22 |
|  | rs2150927 | 0.42 | 20.71 | 0.80 | 4.66 | 0.68 | 0.000 | 0.90 |
| SKIPOGH (night) | rs2486758 | 0.23 | 11.09 | 0.80 | 11.03 | 0.05 | 0.004 | 0.05 |
|  | rs138009835 | 0.10 | -29.08 | 0.64 | -9.60 | 0.24 | -0.006 | 0.06 |
|  | rs2150927 | 0.42 | -0.04 | 0.99 | -1.71 | 0.73 | 0.001 | 0.61 |

$\beta$ = regression coefficient. Analyses were adjusted for age, sex, BMI, time of sampling, and relatedness.

### Associations of morning plasma cortisol and corticosterone with features of metabolic syndrome

In ORCADES (Table 3 and Figure 2A), both higher plasma cortisol and higher plasma corticosterone were associated with lower BMI. After adjustment for BMI, age and sex, both higher plasma cortisol and corticosterone were associated with higher fasting plasma glucose and higher triglycerides, although the magnitude of this association was greater for cortisol than corticosterone. Moreover, higher cortisol, but not corticosterone, was associated with raised systolic blood pressure. Neither cortisol nor corticosterone was significantly associated with total cholesterol or LDL cholesterol. In contrast with these predominant associations with cortisol, lower corticosterone but not cortisol was associated with lower HDL-cholesterol, while higher corticosterone but not cortisol was associated with higher fasting insulin and HOMA-IR. The corticosterone:cortisol ratio was positively associated with fasting glucose, insulin and HOMA-IR but not with BMI, blood pressure or lipid profile.

**Figure 2.**
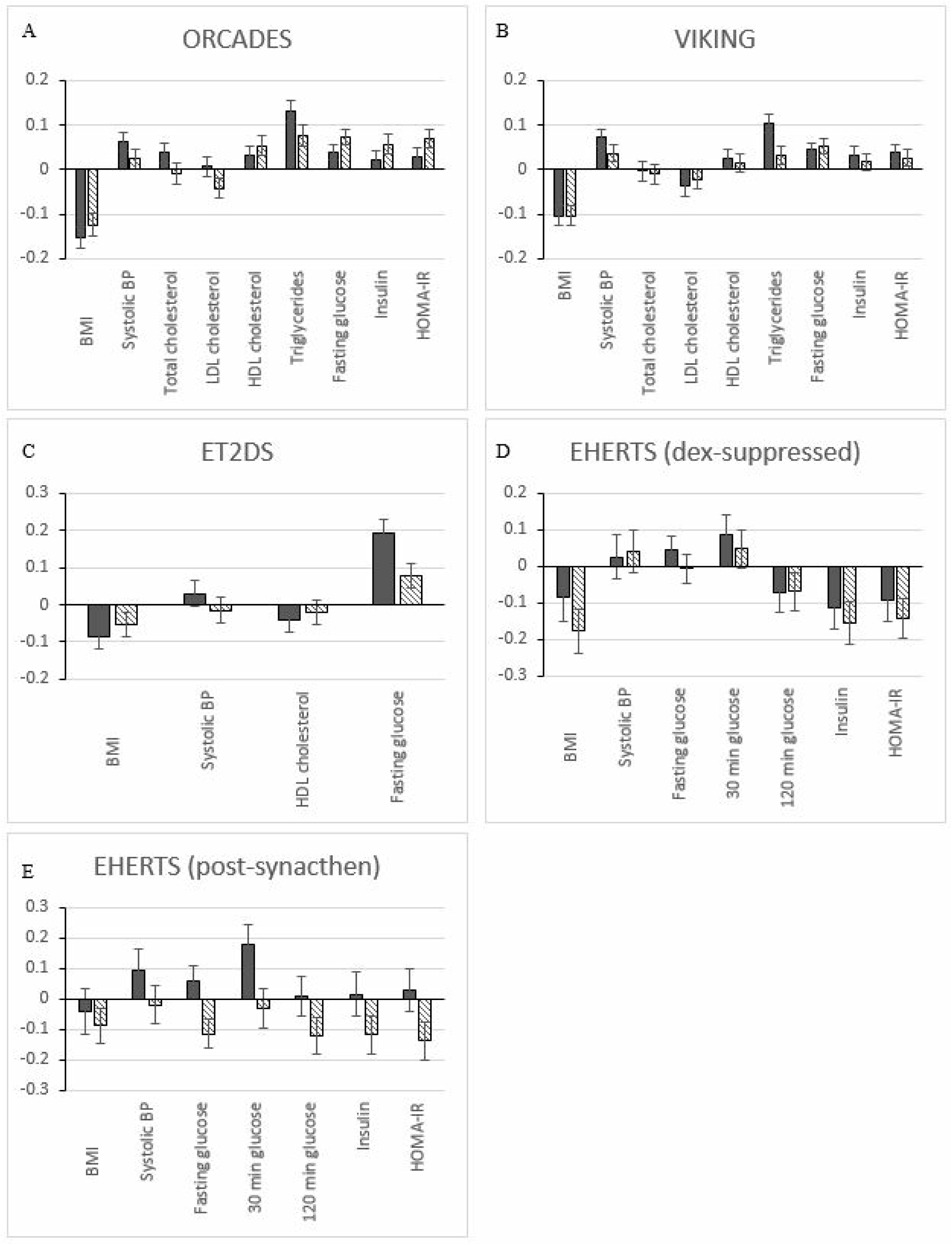
Associations of plasma cortisol and corticosterone with features of metabolic syndrome. Data in A-E are for plasma cortisol (solid bars) and corticosterone (striped bars) from A) Orkney complex diseases study (ORCADES), B) Viking Health Study Shetland (VIKING), C) Edinburgh Type 2 Diabetes Study (ET2DS), D) East Hertfordshire study (EHERTS) after overnight dexamethasone (250 µg) suppression, and E) EHERTS 30 mins after ACTH_1-24_ (1µg). The plots show the SD change (and standard error) in log transformed outcome for a 1SD higher log transformed cortisol or corticosterone.

**Table 3.** Associations of plasma cortisol and corticosterone with features of the metabolic syndrome

|  |  | Cortisol |  |  |  | Corticosterone |  |  |  | Corticosterone:cortisol ratio |  |  |  |
| --- | --- | --- | --- | --- | --- | --- | --- | --- | --- | --- | --- | --- | --- |
|  |  | Unadjusted |  | Adjusted |  | Unadjusted |  | Adjusted |  | Unadjusted |  | Adjusted |  |
| | | $\beta$ | P | $\beta$ | p | $\beta$ | p | B | p | $\beta$ | p | $\beta$ | p |
| ORCADES<br>(n=2018) | BMI | -0.069 | <0.001 | -0.064 | <0.001 | -0.026 | <0.001 | -0.028 | <0.001 | -0.006 | 0.33 | -0.009 | 0.11 |
|  | Systolic BP | -0.021 | 0.01 | 0.022 | 0.001 | -0.004 | 0.41 | 0.005 | 0.25 | 0.003 | 0.48 | -0.002 | 0.57 |
|  | Total cholesterol | -0.006 | 0.61 | 0.021 | 0.09 | -0.002 | 0.73 | -0.002 | 0.73 | -0.010 | 0.17 | -0.008 | 0.27 |
|  | LDL cholesterol | -0.042 | 0.02 | 0.006 | 0.74 | -0.030 | 0.004 | -0.018 | 0.08 | -0.021 | 0.06 | -0.018 | 0.09 |
|  | HDL cholesterol | 0.063 | <0.001 | 0.022 | 0.14 | 0.026 | 0.004 | 0.021 | 0.02 | 0.007 | 0.45 | 0.014 | 0.12 |
|  | Triglycerides | 0.047 | 0.12 | 0.176 | <0.001 | 0.016 | 0.35 | 0.057 | 0.001 | 0.012 | 0.52 | 0.010 | 0.56 |
|  | Fasting glucose | -0.007 | 0.31 | 0.014 | 0.02 | 0.011 | 0.003 | 0.014 | <0.001 | 0.015 | <0.001 | 0.011 | 0.004 |
|  | Insulin | -0.066 | 0.03 | 0.030 | 0.27 | 0.009 | 0.58 | 0.042 | 0.007 | 0.038 | 0.03 | 0.038 | 0.02 |
|  | HOMA-IR | -0.073 | 0.03 | 0.045 | 0.13 | 0.020 | 0.27 | 0.057 | 0.001 | 0.053 | 0.01 | 0.048 | 0.01 |
| VIKING<br>(n=2093) | BMI | -0.047 | <0.001 | -0.035 | <0.001 | -0.018 | <0.001 | -0.014 | <0.001 | -0.012 | 0.001 | -0.010 | 0.004 |
|  | Systolic BP | -0.009 | 0.11 | 0.020 | <0.001 | -0.003 | 0.30 | 0.004 | 0.04 | -0.002 | 0.49 | 0.001 | 0.80 |
|  | Total cholesterol | -0.019 | 0.03 | -0.001 | 0.89 | -0.006 | 0.11 | -0.002 | 0.64 | -0.005 | 0.27 | -0.003 | 0.46 |
|  | LDL cholesterol | -0.048 | <0.001 | -0.021 | 0.09 | -0.013 | 0.02 | -0.005 | 0.35 | -0.006 | 0.32 | -0.002 | 0.69 |
|  | HDL cholesterol | 0.035 | 0.002 | 0.013 | 0.21 | 0.014 | 0.004 | 0.003 | 0.41 | 0.007 | 0.18 | -0.001 | 0.91 |
|  | Triglycerides | 0.017 | 0.48 | 0.112 | <0.001 | -0.015 | 0.13 | 0.014 | 0.13 | -0.022 | 0.04 | -0.006 | 0.59 |
|  | Fasting glucose | -0.006 | 0.22 | 0.012 | 0.008 | 0.001 | 0.72 | 0.006 | 0.002 | 0.002 | 0.32 | 0.005 | 0.02 |
|  | Insulin | -0.041 | 0.10 | 0.037 | 0.09 | -0.020 | 0.06 | 0.008 | 0.37 | -0.017 | 0.15 | 0.001 | 0.96 |
|  | HOMA-IR | -0.047 | 0.08 | 0.048 | 0.04 | -0.020 | 0.09 | 0.014 | 0.15 | -0.015 | 0.25 | 0.005 | 0.62 |
| ET2DS<br>(n=903) | BMI | -0.064 | 0.003 | -0.055 | 0.01 | -0.018 | 0.05 | -0.013 | 0.12 | -0.010 | 0.30 | -0.006 | 0.50 |
|  | Systolic BP | 0.015 | 0.34 | 0.014 | 0.37 | -0.003 | 0.66 | -0.003 | 0.66 | -0.005 | 0.40 | -0.005 | 0.42 |
|  | Fasting glucose | 0.179 | <0.001 | 0.190 | <0.001 | 0.027 | 0.03 | 0.030 | 0.02 | 0.001 | 0.95 | 0.003 | 0.84 |
| EHERTS dex<br>suppressed<br>(n=279) | BMI | -0.018 | 0.13 | -0.017 | 0.18 | -0.036 | 0.003 | -0.036 | 0.004 | -0.023 | 0.14 | -0.024 | 0.12 |
|  | Systolic BP | 0.008 | 0.49 | 0.005 | 0.67 | 0.008 | 0.49 | 0.008 | 0.49 | -0.002 | 0.89 | 0.000 | 1.00 |
|  | Fasting glucose | 0.018 | 0.06 | 0.010 | 0.28 | 0.003 | 0.74 | -0.001 | 0.89 | -0.019 | 0.12 | -0.012 | 0.30 |
|  | 30 min glucose | 0.033 | 0.05 | 0.027 | 0.12 | 0.015 | 0.39 | 0.015 | 0.38 | -0.027 | 0.23 | -0.014 | 0.51 |
|  | 120 min glucose | -0.049 | 0.04 | -0.034 | 0.17 | -0.050 | 0.04 | -0.032 | 0.20 | 0.016 | 0.61 | 0.012 | 0.69 |
|  | Insulin | -0.138 | 0.01 | -0.096 | 0.06 | -0.192 | <0.001 | -0.131 | 0.01 | -0.046 | 0.80 | -0.040 | 0.52 |
|  | HOMA-IR | -0.119 | 0.03 | -0.085 | 0.11 | -0.189 | 0.001 | -0.133 | 0.01 | -0.105 | 0.59 | -0.053 | 0.42 |
| EHERTS<br>after<br>ACTH<sub>1-24</sub><br>(n=279) | BMI | 0.012 | 0.71 | -0.020 | 0.61 | -0.015 | 0.19 | -0.018 | 0.12 | -0.019 | 0.11 | -0.017 | 0.15 |
|  | Systolic BP | 0.014 | 0.65 | 0.048 | 0.19 | -0.007 | 0.58 | -0.004 | 0.77 | -0.014 | 0.32 | -0.011 | 0.41 |
|  | Fasting glucose | -0.031 | 0.19 | 0.037 | 0.20 | -0.033 | 0.003 | -0.027 | 0.01 | -0.021 | 0.06 | -0.026 | 0.02 |
|  | 30 min glucose | 0.019 | 0.66 | 0.146 | 0.01 | -0.023 | 0.25 | -0.010 | 0.63 | -0.021 | 0.32 | -0.019 | 0.35 |
|  | 120 min glucose | 0.092 | 0.15 | 0.010 | 0.90 | -0.054 | 0.06 | -0.056 | 0.05 | -0.072 | 0.02 | -0.059 | 0.05 |
|  | Insulin | 0.136 | 0.32 | 0.036 | 0.82 | -0.120 | 0.04 | -0.099 | 0.07 | -0.046 | 0.02 | -0.096 | 0.09 |
|  | HOMA-IR | 0.103 | 0.48 | 0.072 | 0.66 | -0.153 | 0.01 | -0.126 | 0.03 | -0.485 | 0.02 | -0.123 | 0.04 |

$\beta$ = regression coefficient. P values refer to regression co-efficient, unadjusted or adjusted in multiple regression analyses for co-variables (age, sex, time of sample, relevant medication, and BMI (unless BMI is the outcome variable)). All continuous variables were normalised by log transformation.

In VIKING (Table 3 and Figure 2B), similar associations were observed as in ORCADES. Both cortisol and corticosterone were negatively associated with BMI. Cortisol was more strongly positively associated with systolic blood pressure and triglycerides than corticosterone. The corticosterone:cortisol ratio was negatively associated with BMI and positively associated with fasting glucose.

In ET2DS (Table 3 and Figure 2C), fewer biochemical phenotypes were available, but both higher plasma cortisol and plasma corticosterone were associated with lower BMI and higher fasting plasma glucose, as in ORCADES. However, cortisol and corticosterone were unrelated to systolic blood pressure in ET2DS, and the corticosterone:cortisol ratio was not significantly associated with BMI, blood pressure or glucose.

### Associations of ACTH-stimulated cortisol and corticosterone with features of metabolic syndrome

Given the limited associations of the corticosterone:cortisol ratio in morning plasma samples with features of metabolic syndrome, despite its robust association with *CYP17A1* genotype, we sought to test the association of 17α-hydroxylase activity with features of metabolic syndrome using potentially more sensitive dynamic testing in the EHERTS cohort. See Table 3 and Figures 2D and 2E. After low dose overnight dexamethasone suppression followed by low dose ACTH_1-24_ administration, plasma cortisol and corticosterone levels changed as expected (Table 1). Plasma cortisol was not associated with any features of metabolic syndrome, except that ACTH-stimulated values were positively correlated with plasma glucose 30 min after an oral glucose load. However, plasma corticosterone showed different associations with metabolic syndrome variables. After dexamethasone suppression, lower plasma corticosterone was associated with higher BMI and, independently, with higher fasting insulin and HOMA-IR. After ACTH stimulation, lower plasma corticosterone (and corticosterone:cortisol ratio) was associated with higher plasma glucose before and after an oral glucose load, and with higher HOMA-IR. There were no significant associations between cortisol, corticosterone and blood pressure in the EHERTS cohort.

### Associations of urinary cortisol and corticosterone metabolites with features of metabolic syndrome

In SKIPOGH (Table 4 and Figure 3), after adjustment for confounders (age, sex, study center, relevant medication and BMI), higher cortisol metabolite excretion during daytime and overnight was associated with higher BMI, during the day with higher LDL cholesterol and lower HDL cholesterol, and during the night with higher fasting glucose. Plasma triglycerides were, surprisingly, inversely associated with day and night urinary cortisol metabolite excretion. By contrast, corticosterone metabolite excretion during the day was inversely associated with insulin levels and plasma triglycerides, but during the night was positively associated with fasting glucose and inversely with plasma triglycerides. A lower ratio of corticosterone:cortisol metabolites during both day and night was strongly associated with higher BMI and LDL cholesterol, and during the night was associated with higher systolic BP, higher fasting triglycerides and insulin as well as lower HDL cholesterol levels.

**Figure 3.**
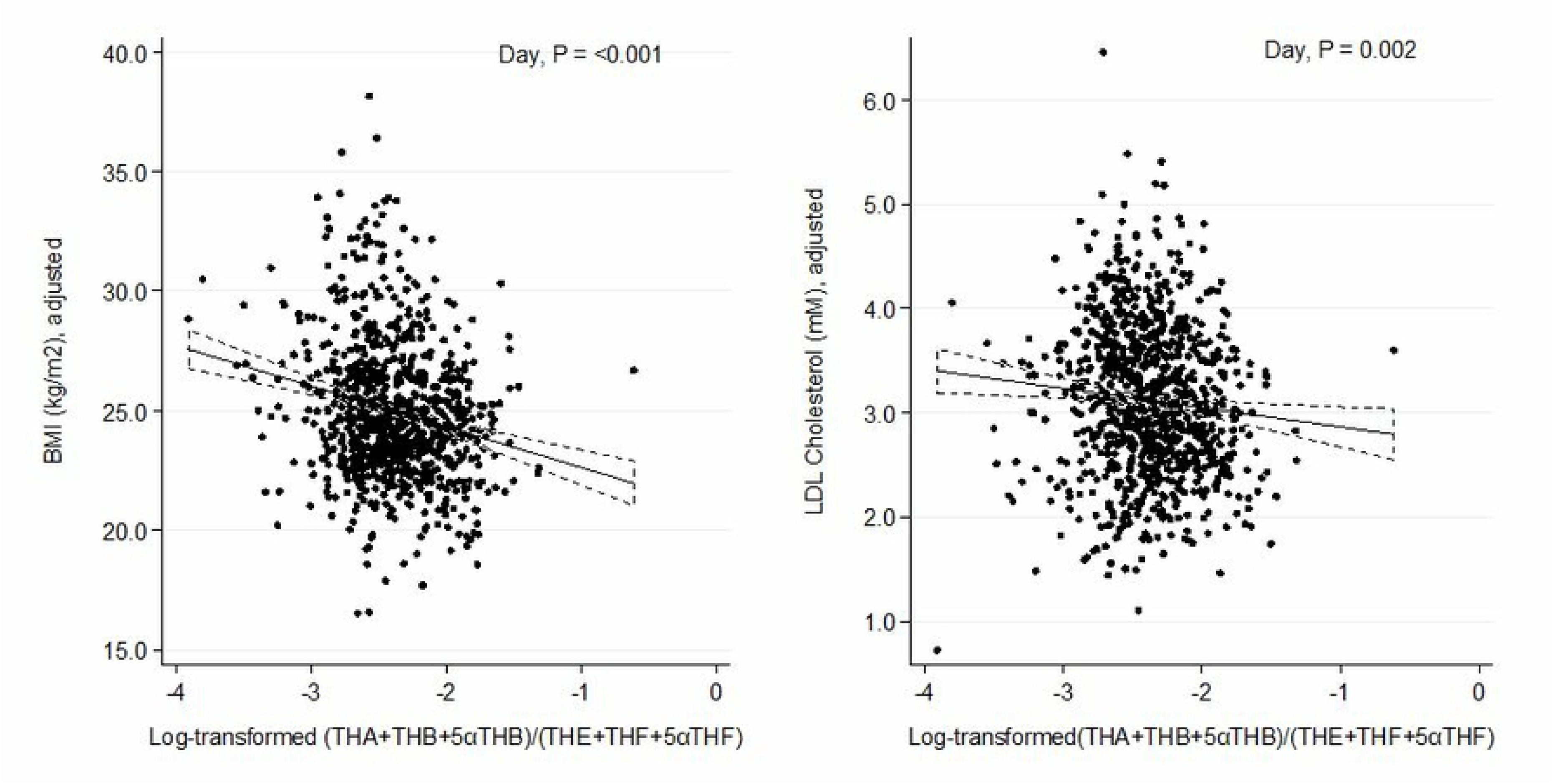
Associations of urine cortisol and corticosterone metabolites with features of metabolic syndrome. Data are from 888 participants in the Swiss Kidney Project on Genes in Hypertension (SKIPOGH). Results are from multiple regression analysis of daytime log-transformed ratio (THA+THB+5αTHB)/(THE+THF+5αTHF) with BMI (left panel) and plasma LDL cholesterol (right panel). The model was adjusted for age, sex, center, lipid- and glucose-lowering and antihypertensive treatment, BMI (for LDL cholesterol only), estimated GFR, urine flow rate, urinary sodium, potassium and creatinine excretion. Results were similar for night-time corticosteroid excretion.

**Table 4.**
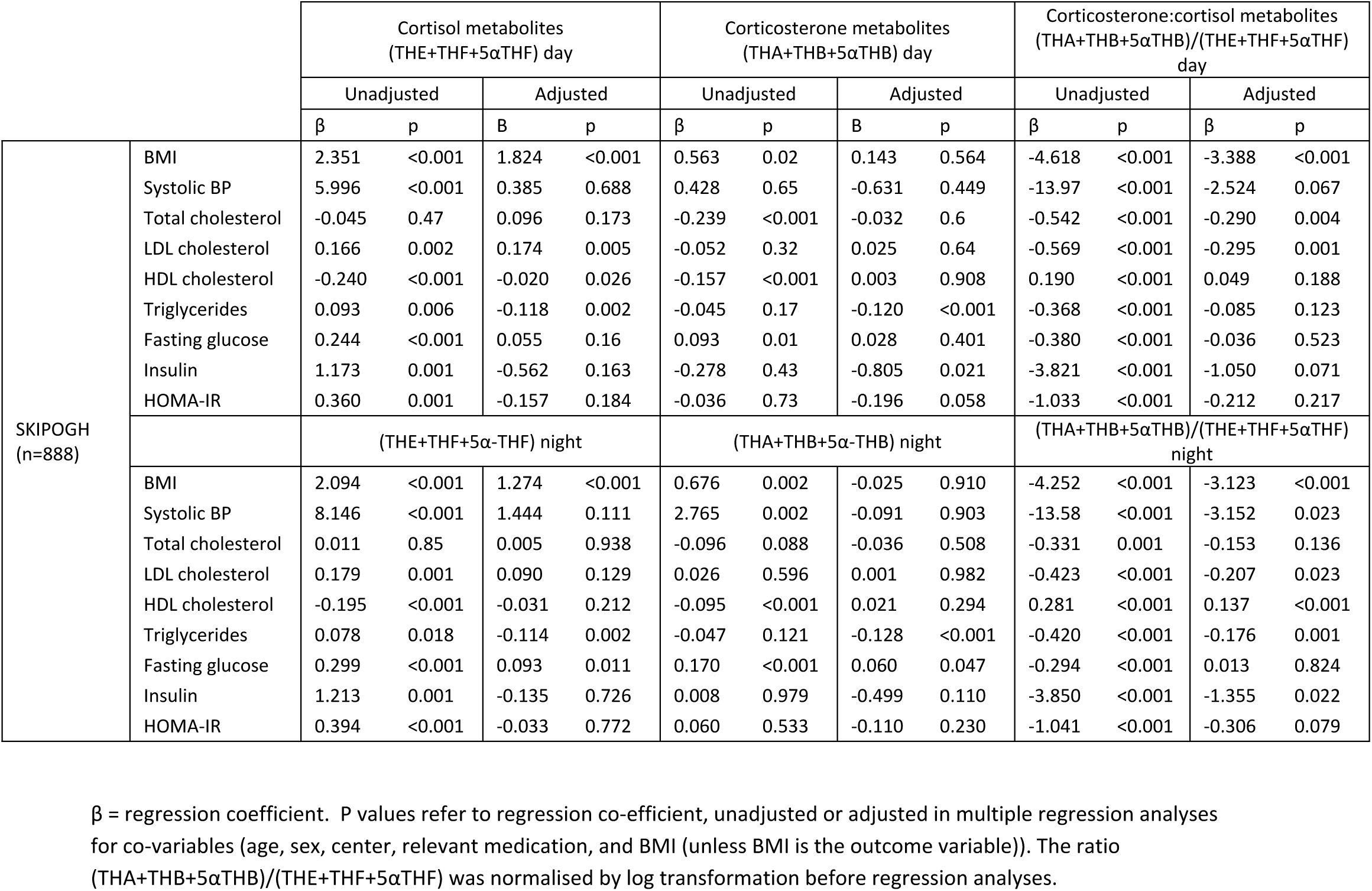
Associations of day- and night-time excretion of urinary cortisol and corticosterone metabolites with features of the metabolic syndrome

| | | Cortisol metabolites<br>(THE+THF+5αTHF) day | | | | Corticosterone metabolites<br>(THA+THB+5αTHB) day | | | | Corticosterone:cortisol metabolites<br>(THA+THB+5αTHB)/(THE+THF+5αTHF)<br>day | | | |
| --- | --- | --- | --- | --- | --- | --- | --- | --- | --- | --- | --- | --- | --- |
|  |  | Unadjusted |  | Adjusted |  | Unadjusted |  | Adjusted |  | Unadjusted |  | Adjusted |  |
| | | $\beta$ | p | B | p | $\beta$ | p | B | p | $\beta$ | p | $\beta$ | p |
| SKIPOGH<br>(n=888) | BMI | 2.351 | <0.001 | 1.824 | <0.001 | 0.563 | 0.02 | 0.143 | 0.564 | -4.618 | <0.001 | -3.388 | <0.001 |
|  | Systolic BP | 5.996 | <0.001 | 0.385 | 0.688 | 0.428 | 0.65 | -0.631 | 0.449 | -13.97 | <0.001 | -2.524 | 0.067 |
|  | Total cholesterol | -0.045 | 0.47 | 0.096 | 0.173 | -0.239 | <0.001 | -0.032 | 0.6 | -0.542 | <0.001 | -0.290 | 0.004 |
|  | LDL cholesterol | 0.166 | 0.002 | 0.174 | 0.005 | -0.052 | 0.32 | 0.025 | 0.64 | -0.569 | <0.001 | -0.295 | 0.001 |
|  | HDL cholesterol | -0.240 | <0.001 | -0.020 | 0.026 | -0.157 | <0.001 | 0.003 | 0.908 | 0.190 | <0.001 | 0.049 | 0.188 |
|  | Triglycerides | 0.093 | 0.006 | -0.118 | 0.002 | -0.045 | 0.17 | -0.120 | <0.001 | -0.368 | <0.001 | -0.085 | 0.123 |
|  | Fasting glucose | 0.244 | <0.001 | 0.055 | 0.16 | 0.093 | 0.01 | 0.028 | 0.401 | -0.380 | <0.001 | -0.036 | 0.523 |
|  | Insulin | 1.173 | 0.001 | -0.562 | 0.163 | -0.278 | 0.43 | -0.805 | 0.021 | -3.821 | <0.001 | -1.050 | 0.071 |
|  | HOMA-IR | 0.360 | 0.001 | -0.157 | 0.184 | -0.036 | 0.73 | -0.196 | 0.058 | -1.033 | <0.001 | -0.212 | 0.217 |
| | | (THE+THF+5α-THF) night | | | | (THA+THB+5α-THB) night | | | | (THA+THB+5αTHB)/(THE+THF+5αTHF)<br>night | | | |
|  | BMI | 2.094 | <0.001 | 1.274 | <0.001 | 0.676 | 0.002 | -0.025 | 0.910 | -4.252 | <0.001 | -3.123 | <0.001 |
|  | Systolic BP | 8.146 | <0.001 | 1.444 | 0.111 | 2.765 | 0.002 | -0.091 | 0.903 | -13.58 | <0.001 | -3.152 | 0.023 |
|  | Total cholesterol | 0.011 | 0.85 | 0.005 | 0.938 | -0.096 | 0.088 | -0.036 | 0.508 | -0.331 | 0.001 | -0.153 | 0.136 |
|  | LDL cholesterol | 0.179 | 0.001 | 0.090 | 0.129 | 0.026 | 0.596 | 0.001 | 0.982 | -0.423 | <0.001 | -0.207 | 0.023 |
|  | HDL cholesterol | -0.195 | <0.001 | -0.031 | 0.212 | -0.095 | <0.001 | 0.021 | 0.294 | 0.281 | <0.001 | 0.137 | <0.001 |
|  | Triglycerides | 0.078 | 0.018 | -0.114 | 0.002 | -0.047 | 0.121 | -0.128 | <0.001 | -0.420 | <0.001 | -0.176 | 0.001 |
|  | Fasting glucose | 0.299 | <0.001 | 0.093 | 0.011 | 0.170 | <0.001 | 0.060 | 0.047 | -0.294 | <0.001 | 0.013 | 0.824 |
|  | Insulin | 1.213 | 0.001 | -0.135 | 0.726 | 0.008 | 0.979 | -0.499 | 0.110 | -3.850 | <0.001 | -1.355 | 0.022 |
|  | HOMA-IR | 0.394 | <0.001 | -0.033 | 0.772 | 0.060 | 0.533 | -0.110 | 0.230 | -1.041 | <0.001 | -0.306 | 0.079 |

$\beta$ = regression coefficient. P values refer to regression co-efficient, unadjusted or adjusted in multiple regression analyses for co-variables (age, sex, center, relevant medication, and BMI (unless BMI is the outcome variable)). The ratio (THA+THB+5αTHB)/(THE+THF+5αTHF) was normalised by log transformation before regression analyses.

## DISCUSSION

This is the largest study to date investigating associations between endogenous glucocorticoids and features of metabolic syndrome. Using 3 cohorts comprising 3200 participants, we confirm previous observations (13, 31–34) that elevated morning plasma cortisol is associated with higher blood pressure and blood glucose in subjects with and without type 2 diabetes. Moreover, we confirm previous observations (35) in by far the largest sample to date that urinary cortisol metabolite excretion is increased in association with obesity and dyslipidemia. With a large number of participants, the chance of statistical errors is much reduced.

This is the first study to investigate whether similar associations exist for corticosterone. We measured plasma corticosterone and analysed variability in *CYP17A1* to determine whether ‘relative corticosterone deficiency’, favouring production of cortisol over corticosterone, underlies associations between variation in *CYP17A1* and features of metabolic syndrome identified in GWAS consortia. The sample size was too small to detect associations at genome-wide significance, but using a candidate gene approach we demonstrated that variation in *CYP17A1* associates with plasma corticosterone:cortisol ratio in a general adult population, and confirmed our findings in two additional cohorts (one in a general adult population and one in a cohort of patients with type 2 diabetes). Moreover, these observations were supported by an association of *CYP17A1* genotype with urinary corticosterone/cortisol metabolite ratios. The association was restricted to just one SNP of 3 that are known to have functional effects on transcription *in vitro* (27), rs2486758; the replication in a second independent cohort, however, makes it highly unlikely to be due to chance. rs2486758 is located in the 5’ regulatory region of *CYP17A1* and in LD with rs2150927 but less so with rs138009835 or the variants previously associated with hypertension by GWAS (27). Paradoxically, the major T allele of rs2486758, which we find associated with ‘relative corticosterone deficiency’, induced lower, rather than higher, CYP17A1 expression *in vitro* (27) but this may be an artefact of the *in vitro* system or rs2486758 may be linked with other functional variants.

Previous large GWAS consortia have shown opposite directions of association between variants in *CYP17A1* with hypertension versus other features of metabolic syndrome. We therefore anticipated that ‘relative corticosterone excess’ is associated with higher blood pressure while ‘relative corticosterone deficiency’ is associated with cortisol-mediated insulin resistance and obesity. However, despite elevated corticosterone excretion rate being reported previously in hypertension (36), we did not find an association between morning plasma corticosterone and blood pressure. Moreover, our data show that morning plasma cortisol and corticosterone were positively correlated. Further, plasma cortisol and corticosterone showed similar associations with BMI; it is likely that reduced plasma corticosterone in obesity is explained by increased activity of A-ring reductases which also underlie increased clearance of cortisol in obesity (37). When we did find discrepancies in associations with plasma cortisol or corticosterone, it was generally higher, rather than lower, morning plasma corticosterone that was the stronger predictor than morning plasma cortisol, e.g. of fasting plasma glucose, plasma insulin and HOMA-IR. This suggests that *CYP17A1* genotype is not the major determinant of morning plasma corticosterone:cortisol ratio. It is possible that insulin resistance or deficiency causes a shift in steroidogenesis in favour of corticosterone rather than cortisol, since *CYP17A1* expression is up-regulated by insulin (38). Alternatively, the combined elevation of cortisol and corticosterone with insulin resistance could reflect central activation of the HPA axis, consistent with altered responses to habituation and stress (34).

Associations between morning plasma glucocorticoid levels and lipid profile provided limited support for our hypothesis of ‘relative corticosterone deficiency’. Thus, elevated triglycerides were consistently associated with plasma cortisol and less strongly with plasma corticosterone, and low HDL-cholesterol was associated only with low corticosterone, although the latter association has been reported previously for cortisol (39). However, effects of glucocorticoids on lipid metabolism are complex (40), and previous studies investigating associations of glucocorticoid excess with dyslipidaemia have been inconsistent (13).

A different pattern emerged when we studied associations of corticosterone and cortisol in EHERTS subjects who had received dexamethasone to suppress any compensatory increase or primary drive to the HPA axis, and a fixed dose of ACTH_1-24_ to stimulate cortisol and corticosterone. The results are striking, with opposite associations of cortisol and corticosterone with metabolic syndrome features, consistent with a primary shift in adrenocortical production in favour of cortisol rather than corticosterone in metabolic syndrome. Since, as a result of the activity of ABCB1 excluding cortisol from the brain (15), corticosterone is thought to make a disproportionate contribution to negative feedback suppression of the HPA axis in humans, these observations could provide a key insight into the activation of the HPA axis and elevated plasma cortisol in metabolic syndrome. This insight is obscured by measurement of morning plasma corticosterone and cortisol alone, perhaps because of greater variability as a result of unmeasured confounding.

Fasting plasma corticosterone was low in the ET2DS cohort, whereas cortisol was similar to the values in the ORCADES cohort (mean corticosterone:cortisol was 3.0% in ET2DS vs. 7.5% in ORCADES, p<0.0001). This is unlikely to be due to older age of the ET2DS cohort, since corticosterone:cortisol ratio was not associated with age (r^2^ = 0.01, p=0.27). Again, insulin deficiency might favour cortisol rather than corticosterone secretion, consistent with loss of up-regulation of *CYP17A1* by insulin (38). If confirmed, this mechanism may exacerbate the ‘relative corticosterone deficiency’ apparent with metabolic syndrome in EHERTS, and thereby exacerbate HPA axis activation in type 2 diabetes.

Given the limitations of morning plasma cortisol and corticosterone, we investigated the hypothesis of relative corticosterone deficiency in another cohort, the SKIPOGH study, in which integrated measurements of cortisol and corticosterone metabolite excretion in urine were available in a large number of participants. These results confirmed that the pattern observed for plasma glucocorticoids after ACTH_1-24_ stimulation in EHERTS, with relatively low corticosterone and high cortisol, is similarly associated in urine with glucose and lipid metabolism and, strikingly, with obesity. Surprisingly, ‘relative corticosterone deficiency’ was also associated with higher systolic clinic blood pressure, although previous studies in this cohort have suggested that ambulatory blood pressure is inversely associated with CYP17A1 activity only in the presence of high sodium intake (8).

An important limitation of these studies is the cross-sectional design which precludes inferences of causation. Moreover, genotype data was not available for EHERTS subjects because samples of DNA from this cohort have been exhausted. Indeed, no sample of sufficient size exists in which ACTH stimulation tests have been conducted to be adequately powered to detect associations of steroidogenic response with *CYP17A1* genotype. Also, effects of intensive lipid-lowering and antihypertensive treatments in the ET2DS may have obscured associations with corticosterone and cortisol. A further limitation is that the dynamic HPA axis tests and urinary steroid analyses were conducted in different cohorts from the measurement of fasting plasma glucocorticoids. The corollary is, however, that we replicated associations of candidate *CYP17A1* genotype with phenotype in three independent studies and we have been drawn upon comprehensive datasets to adjust for a number of potentially confounding variables and assess the corticosterone:cortisol relationship in urine as well as plasma. We present our findings unadjusted for multiple testing because, as discussed above, there are distinct associations of glucocorticoid levels with individual components of metabolic syndrome such that each analysis tests a distinct hypothesis.

In conclusion, elevated morning plasma corticosterone accompanies insulin resistance and elevated cortisol in metabolic syndrome, consistent with activation of the HPA axis at the time of blood sampling. However, in addition, there are marked discrepancies between associations with ACTH-stimulated cortisol and corticosterone with metabolic syndrome which likely reflect genetically determined differences in adrenal steroidogenesis, particularly by 17α-hydroxylase, and may also be influenced by dysregulated insulin signaling. When sustained HPA axis activation is assessed by urinary metabolites, the variation in adrenal steroidogenesis predominates in the associations with metabolic syndrome and, in particular, obesity. These findings throw the spotlight on corticosterone and suggest that further dissection of its biology in humans may be fruitful in understanding the basis for altered glucocorticoid signaling in metabolic syndrome.

## ACKNOWLEDGEMENTS

We are grateful to: Lynne Ramage and Jill Harrison for technical support; Stela McLachlan for data processing in ET2DS; the study nurses Marie-Odile Levy, Guler Gök-Sogüt, Ulla Schüpbach, Dominique Siminski and Sandrine Estoppey for the data collection in SKIPOGH; and all of the participants and researchers involved. We also acknowledge the investigators in the Swiss Kidney Project on Genes in Hypertension (SKIPOGH) team: (in alphabetical order) Daniel Ackermann^1^, Murielle Bochud^2^, Michel Burnier^3^, Georg Ehret^4^, Idris Guessous^5^, Pierre-Yves Martin^6^, Markus Mohaupt^1^, Fred Paccaud^2^, Antoinette Pechère-Bertschi^7^, Belen Ponte^6^, Menno Pruijm^3^, and Bruno Vogt^1^.

^1^University Clinic for Nephrology, Hypertension and Clinical Pharmacology, Inselspital, Bern University Hospital, University of Bern, Switzerland

^2^Institute of Social and Preventive Medicine (IUMSP), Lausanne University Hospital and University of Lausanne, Switzerland

^3^Department of Nephrology, Lausanne University Hospital, Lausanne, Switzerland.

^4^Cardiology, Department of Specialties of Internal Medicine, Geneva University Hospitals, Geneva, Switzerland.

^5^Unit of Population Epidemiology, Department of Community Medicine and Primary Care and Emergency Medicine, Geneva University Hospitals, Geneva, Switzerland.

^6^Department of Nephrology, Geneva University Hospitals, Geneva, Switzerland.

^7^Unit of Hypertension, Departments of Community Medicine and Primary Care and Emergency Medicine, Geneva University Hospitals, Switzerland.

